# Automated behavioral profiling using neural networks reveals differences in stress-like behavior between cave and surface-dwelling *Astyanax mexicanus*

**DOI:** 10.1101/2025.01.30.635725

**Authors:** Naresh Padmanaban, Rianna Ambosie, Stefan Choy, Shoshanah Marcus, Simon R.O. Nilsson, Alex C. Keene, Johanna E. Kowalko, Erik R. Duboué

## Abstract

Behavioral stress responses allow animals to quickly adapt to local environments and are critical for survival. Stress responses provide an ideal model for investigating the evolution of complex behaviors due to their conservation across species, critical role in survival, and integration of behavioral and physiological components. The Mexican cavefish (*Astyanax mexicanus*) has evolved dramatically different stress responses compared to river-dwelling surface fish morphs, providing a model to investigate the neural and evolutionary basis of stress-like responses. Surface morphs inhabit predator-rich environments whereas cave-dwelling morphs occupy predator-free habitats. While these key ecological variables may underlie differences in stress responses, the complexity of the behavioral differences has not been thoroughly examined. By leveraging automated pose-tracking and machine learning tools, we quantified a range of behaviors associated with stress, including freezing, bottom-dwelling, and hyperactivity, during a novel tank assay. Surface fish exhibited heightened stress responses characterized by prolonged bottom-dwelling and frequent freezing, while cavefish demonstrated reduced stress behaviors, marked by greater exploration and minimal freezing. Analysis of F2 hybrids revealed that a subset of behaviors, freezing and bottom-dwelling, co-segregated, suggesting shared genetic or physiological underpinnings. Our findings illustrate the power of computational tools for high-throughput behavioral phenotyping, enabling precise quantification of complex traits and revealing the genetic and ecological factors driving their evolution. This study provides a framework for understanding how integrated behavioral and physiological traits evolve, offering broader insights into the mechanisms underlying the diversification of animal behavior in natural systems.

## Introduction

Behavioral and physiological responses to stressors are critical for survival and are subject to natural selection across the animal kingdom (Blackledge and Gillespie, 2004; Nesse et al., 2016). For example, both across species and within species, individuals or populations in high-predator environments exhibit heightened stress responses, such as elevated glucocorticoid levels and reduced exploratory behavior, compared to those in predator-free settings (Mateo, 2007; Fischer et al., 2014; Heinen-Kay et al., 2016). These differences underscore the role of ecology in shaping the evolution of stress-related traits (Barton, 2002; Huber et al., 2017; Chin et al., 2018; Chin et al., 2020). While previous studies have provided an understanding of how ecological pressures influence stress responses, much of this work relies on manual scoring of behavior, which is time-consuming and vulnerable to biases and experimental error. To uncover the mechanisms driving evolutionary trajectories of stress responses, it is crucial to adopt approaches that capture the full spectrum of behavioral components and their variation in a standardized manner that does not differ between annotators.

Recent advances in automated analysis of animal behavior, including tracking and machine learning-based approaches, have significantly enhanced our understanding of the biological basis of complex behaviors (Branson et al., 2009; Perez-Escudero et al., 2014; Mathis et al., 2018). For example, pose-detection is capable of tracking multiple body parts of an animal over time rather than a single center point, permitting the estimation of a wide range of behaviors that are undetectable with simplified tracking software (Mathis et al., 2018). Additionally, behavioral classification using deep neural networks enables the automated detection of multiple complex behaviors in an unbiased and high-throughput manner (Goodwin et al., 2024; Goodwin and Golden, 2024). While these tools have been transformative in studying the biological and neural underpinnings of behavior, they have not yet been widely applied to exploring natural variation in behavior across populations and species. Leveraging these innovative tools presents an unprecedented opportunity to investigate how complex behaviors vary and evolve in response to ecological and evolutionary pressures.

The blind Mexican cavefish, *Astyanax mexicanus*, has emerged as a powerful system for examining how morphological and behavioral traits change over evolutionary time. The species consists of two dramatically different morphs: populations of eyed pigmented fish live in above-ground rivers and streams in Mexico and southern Texas, and at least 30 populations of cave-dwelling fish inhabit caves within Northeast Mexico’s Sierra de El Abra and Sierra de Guatemala regions (Mitchell et al., 1977; Jeffery, 2001, 2009; Gross, 2012). Cave dwelling *A. mexicanus* have evolved eye regression and reductions in pigmentation, as well as many behavioral traits including reduced aggression, schooling, and sleep and alterations to feeding (Protas et al., 2006; Protas et al., 2007; Duboué et al., 2011; Aspiras et al., 2015; Jaggard et al., 2018; Lloyd et al., 2018; Pierre et al., 2020; Rodriguez-Morales et al., 2022).

We have identified robust differences between stress-like responses of surface and cavefish populations (Chin et al., 2018; Chin et al., 2020). Following a stressor such as a confined space, surface fish have significantly higher cortisol levels relative to cavefish subjected to the same stimuli (Gallo and Jeffery, 2012). Similarly, we have shown that both larval and adult surface fish have elevated behavioral measures of stress-responses compared to multiple populations of cavefish (Chin et al., 2018; Chin et al., 2020). These findings reveal robust differences in stress response between surface fish and independently evolved populations of cavefish, which, combined with their comparative biology and amenability to genetic manipulation, make *A. mexicanus* a powerful model for identifying mechanisms underlying the evolution of stress responses.

The novel tank test is a widely used assay for examining stress behavior in fish (Levin et al., 2007; Cachat et al., 2010). In this assay, adult fish are placed in an unfamiliar tank and allowed to explore for a 10-min period. Initially, fish swim in the bottom portion of the tank, yet over time as the fish acclimates, it begins to explore top and bottom halves with near-equal frequency. Application of noxious compounds results in fish positioning themselves to the bottom half, whereas pre-treatment with anxiolytics results in fish disproportionately swimming in the top portion, suggesting time spent in the bottom is a measure of stress levels (Bencan et al., 2009; Cachat et al., 2010; Mathuru et al., 2012). We have shown that surface fish prefer the bottom portion of the tank for the majority of the recording period, whereas cavefish quickly explore both halves of the tank (Chin et al., 2018). While time in the bottom is considered a valid measure of stress levels, fish show a myriad of stress-associated behaviors such as freezing and erratic swimming (Schreck et al., 2016), which have not been quantified in *A. mexicanus*.

Here, we sought to identify differences in stress behavior using precise and automated behavioral analysis. We applied pose-tracking to surface and cave morphs of *Astyanax mexicanus* in the novel tank assay. Using a machine learning approach with deep neural networks, we automatically evaluated a range of behaviors associated with stress. Consistent with previous studies, we found that surface fish spent more time at the bottom of the tank compared to their cave-dwelling counterparts. Additionally, other behaviors, such as freezing and hyperactivity, differed quantitatively between the two morphs. High-throughput behavioral analysis of surface × cave hybrid fish revealed that these behaviors clustered into two main groups, suggesting that correlated behavioral traits may have evolved through the same genetic mechanisms as each other. Together, these findings provide a comprehensive assessment of behavior in an evolutionary model and establish a foundation for the automated evaluation of complex traits.

## Methods

### Ethics Statement

All experimental procedures were carried out in accordance with approval from the Institutional Animal Care and Usage Committee (IACUC) at Florida Atlantic University, protocol numbers A17–21 and A15–32 and Lehigh University. All efforts were made to ensure health of the fish, and behavioral procedures were designed to minimize any unnecessary stress or pain.

### Animal Maintenance

All experiments in this study were performed on laboratory-born surface (Texas) fish, cavefish from the Pachón populations and F2 hybrids (Rio Choy x Pachón F2’s). Animal care was conducted as previously described. Briefly, fish were maintained on a custom-designed closed recirculating aquatics system (Aquaneering) in 18-37 L glass tanks with water temperature maintained at 23±1°C. The aquatics facility is housed in a humidity-controlled room and is maintained on a constant 14:10 light:dark cycle with light intensity of 25-40 lux. Fish were fed Ziegler pellets daily, and the diet was periodically supplemented with California Black Worms (Kozol et al., 2023).

### Behavioral Recordings

Behavioral experiments were performed in the novel tank diving test in 18 cm (w) x 11 cm (l) x 14 cm (h) plastic tanks (recording chamber) set in front of custom-designed infrared (IR) light sources (850 nm; Fig. 1A). Before recording, 1.5-2 L of fresh system water was added to each tank. Adult fish were allowed to sit in the room acclimatized for 10 min in individual 500 mL beakers before recording. After the 10 min acclimation period, each fish was introduced into one of the recording chambers. Locomotor activity was recorded along the z-axis using a scientific cMOS camera (Basler aCA 1300-200um or Basler aCA 640-90um) attached to a 16mm f/1.4 (Edmund Optics) fixed focal length lens. The camera was positioned in front of the recording tank such that the z-axis of the fish was monitored. Video records were collected using Pylon Viewer software (Basler). All behavioral recordings were collected between 9:00 am and 6:00 pm. Recordings were conducted for a 10-minute period at 25-30 fps and saved as an .mp4 file. Adult fish were transferred into a holding tank following recording. All fish were transferred back into their home tanks after the experiment ended.

**Fig 1.**
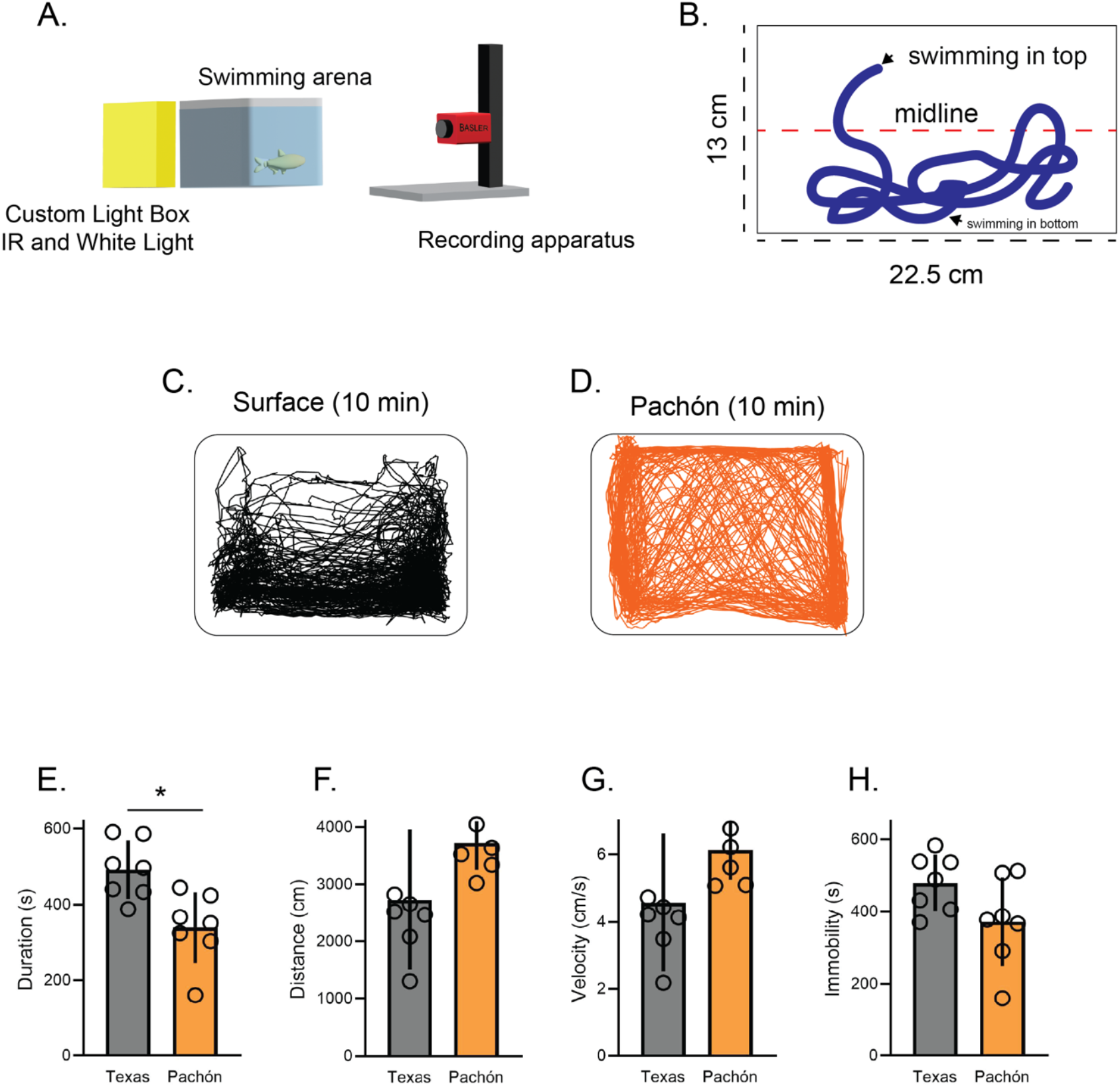
Cave morphs show reduced stress behaviors compared to surface conspecifics. (A) Custom recording setup used to conduct behavioral recordings of A. mexicanus. (B) Schematic of tank dimensions and expected results. Fish typically spend the first few minutes swimming in the bottom portion of the tank, but at the end of the period the fish explore the top and bottom with near equal frequency. Studies have shown that time at the bottom is indicative of stress. (C) The path traveled by a Surface morph during a 10-minute period. (D) The path traveled by a cave morph during a 10-minute period. (E) Quantification of total duration spent in the bottom portion reveals significant differences between surface and cavefish (p = 0.006). (F-H) Quantification of distance traveled, velocity, and immobility trended on significance (p = 0.069; p = 0.089; p = 0.074). E-H quantified using Ethovision XT 13. For surface fish, n=7; for Pachón, n=7. Asterisks indicate significance below p = 0.05.

### Assessing behavior using Ethovision XT 13

A subset of behavior recordings were analyzed using Ethovision XT, as previously described (Chin et al., 2018). Briefly, movies of fish behavior in the novel tank test were imported into Ethovision as .mp4 files. A project tracking ‘zebrafish adult’ as subject type was formed. The environment was set to ‘open field square’ area and method of tracking set to ‘center point tracking.’ Area settings were set with two zones, one top half and the other bottom half, with detection settings to ‘automatic’. Following tracking, time in top and bottom half, total distance traveled, velocity and immobility were calculated from the final tracks.

### DeepLabCut tracking

DeepLabCut (DLC) (Mathis et al., 2018) was used for automated tracking using either a custom-built Desktop computer with a Ryzen 9 3900X 3.8GHz processor (AMD), 64 GB RAM (Team T-force Vulcan 3200 CL16) or a computing device of similar performance. To prepare a training dataset for DLC, we selected 5 body parts (mouth, body_midpoint, dorsal_fin, tail_midpoint, peduncle_base) to be tracked through the duration of each video (see Fig 2). We then manually selected videos and used DLC’s “`extract_frames()”` function to select 20 random frames from each video, then marked the previously indicated body parts in each of these frames using the function “`label_frames()`”. After checking these annotations with DLC’s “`check_labels()`” function, we created a training dataset using DLC’s “`create_training_dataset()`” function. To perform pose-estimation in our videos, we applied DLC’s “`train_network()`” function to this dataset. We then tracked poses in all videos using DLC’s “`analyze_videos()`” function, generating a .csv file describing the x- and y-coordinates of each of the 5 body parts. A total of 15 videos was used to train the model. The quality of the model was evaluated by a trained experimenter comparing the automated annotations to the labeled video records.

**Fig 2.**
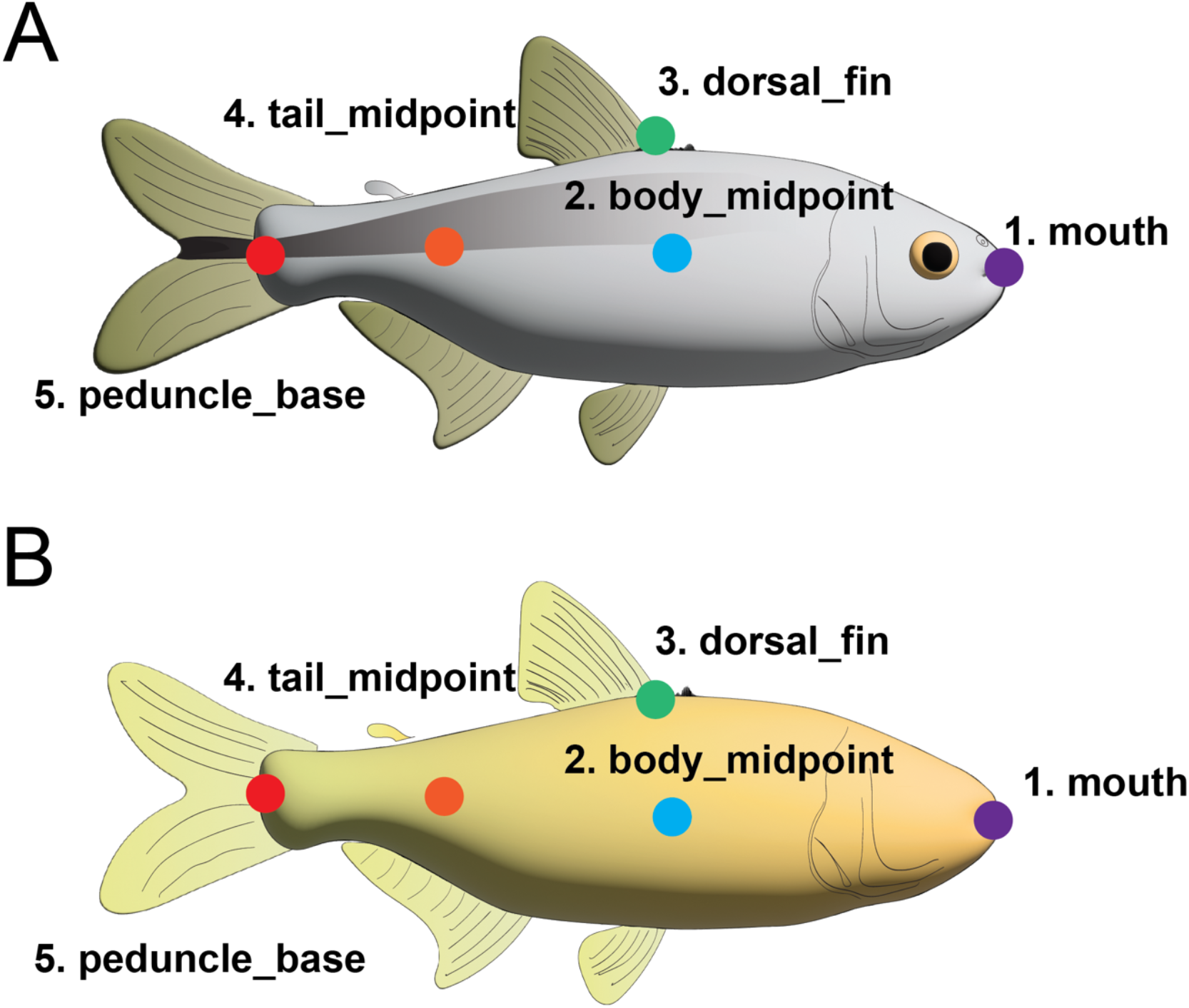
Body part annotations of cave and surface fish using DeepLab Cut. Body part annotations are shown for surface fish (A) and cavefish (B). The body parts spanned the entire fish, including the mouth, midpoint of the body, dorsal fin, midpoint of the tail, and the peduncle base.

### Boris Annotations

To manually record behavioral annotations for our subjects, we used Behavioral Observation Research Interactive Software (BORIS) (Friard and Gamba, 2016). A subset of our videos (n=15) were imported into BORIS to generate model data for use in our behavioral classifier. This subset included video recordings of both surface and cave morphs. A total of nine behaviors were tracked: Normal Swimming, Fast Swimming, Freezing, Normal Turning, Erratic Turning, Floor Skimming, Wall Bumping, Top, and Bottom (see table 1). Videos were inspected frame by frame by a trained experimenter, and the events were specified as state events for each given frame. After completion, data was exported as a .csv file for analysis and an ethogram (.png) to visualize when the behaviors occurred.

**Table 1.**
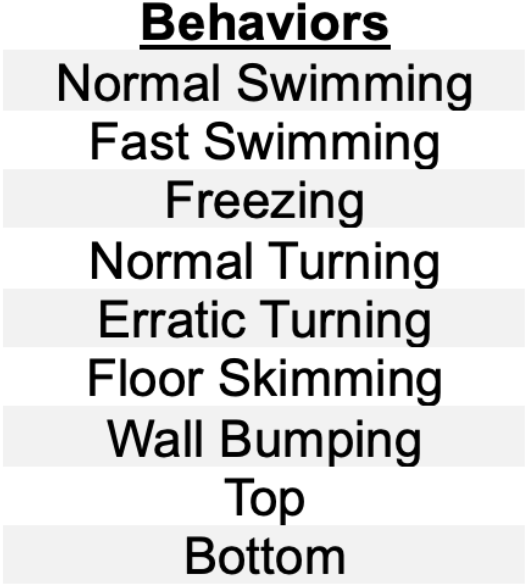
Behaviors assessed through machine learning.

### Behavioral Classifier and Training

Behavioral classifiers were implemented using Simple Behavioral Analysis (SimBA) (Goodwin et al., 2024) on the custom-built computer. A project configuration file was created with 9 classifiers (the behaviors annotated in BORIS). A SimBA-specific file (i.e., body part configuration file) delineating the tracking points that DeepLab Cut used was then created in SimBA. Behavioral recordings (.mp4), DeepLabCut tracking data (.csv), and BORIS annotations (.csv) were imported into the program. The diagonal length of the recording tank was set as the real-world distance for each of the recordings for use in calculating distance traveled velocity. Features were extracted using a custom feature-extraction python script developed by our labs (see supplemental material). Post feature-extraction, a custom python script was used to append the behavior annotations to the extracted features (see supplemental script). To train the classifiers, the following training criteria was used: RF estimators: 2000; max features: sqrt; criterion: gini; test size: 0.2; Train-test split type: frames; Under sample setting: Random under sample; Under sample ratio: 0.15). The classifiers or behaviors of interest were then run for the total number of videos in the dataset for both surface and multiple cave morphs. A threshold was set for each behavior (the level of probability required for a behavior to be recognized as a behavior) and the minimum bout length (minimum amount of time the behavior must occur) for each behavior was set. After analysis, SimBA generated the same videos with the behavioral predictions overlaid. The quality of the model was evaluated by comparing our manual annotated behaviors with that generated by SimBA for the same video.

### Statistics

We used the programming language Python (v3.12.0), as well as the Prism software GraphPad (v10), to perform statistical tests and create graphs. We used unpaired, parametric Student’s t-tests to test for significance between the two morphs in terms of behavior.

## Results

### Cavefish exhibit reduced stress-like behavior

Previous work from our lab suggested that cavefish spend less time bottom-dwelling in the novel tank test relative to their surface conspecifics (Chin et al., 2018). To confirm these results, we assayed surface and Pachón cavefish in the novel tank test (Fig 1A, B). Surface and Pachón fish were placed in an unfamiliar tank and their locomotor activity and tank location was recorded for a period of 10 minutes. We then used Ethovision XT to evaluate the duration of time at the top and bottom. Similar to previous studies, surface fish preferred the bottom of tank (Fig. 1C). In contrast, the Pachón fish explored a significant portion of the top of the tank over the 10-min recording (Fig 1D). Quantification of total duration spent in the bottom revealed significant differences (p=0.006). We also evaluated additional metrics including distance traveled, velocity and immobility. Surface fish exhibited a trend toward traveling shorter distances (p=0.07), swam at slower velocities (p=.09), and had more time immobile (p=0.07) compared to Pachón cavefish (Fig. 1E-H). These trends suggest additional differences in behavior between cave and surface fish in response to the novel tank assay, and highlight the complexity of the behavior.

### Machine learning pipeline to predict stress-like behaviors

Stress induces a range of behaviors, including reduced exploration, freezing, hyperactivity, and erratic turning. However, these behaviors have not been systematically quantified in *A. mexicanus*. To identify distinct behavioral components associated with stress, we implemented a machine learning pipeline incorporating a behavioral classifier to identify and quantify nine distinct behaviors relevant to stress (Table 1). Fish behavior during the novel tank assay was first recorded, and body parts of fish in each record was tracked using DeepLab Cut. We tracked body parts throughout the fish, including the mouth, midpoint of the body, dorsal fin, mid-point of the tail, and peduncle base (Fig 2A,B). Next a subset (n=15) of videos which were to be used to train the behavioral classifier (training videos) were manually analyzed by a trained experimenter using Behavioral Observation Research Interactive Software (BORIS). BORIS allows frame-by-frame classification of behaviors, generating a comprehensive dataset of annotated behaviors. Each of the nine behaviors in Table 1 were annotated in each of the 15 training videos. To train SimBA, a machine learning behavioral classifier, pose-tracking data from DeepLab Cut and the manually scored BORIS annotations were inputted into the program, which enabled the automated classification of behaviors across the full entire recording period (Fig. 3A).

**Fig 3.**
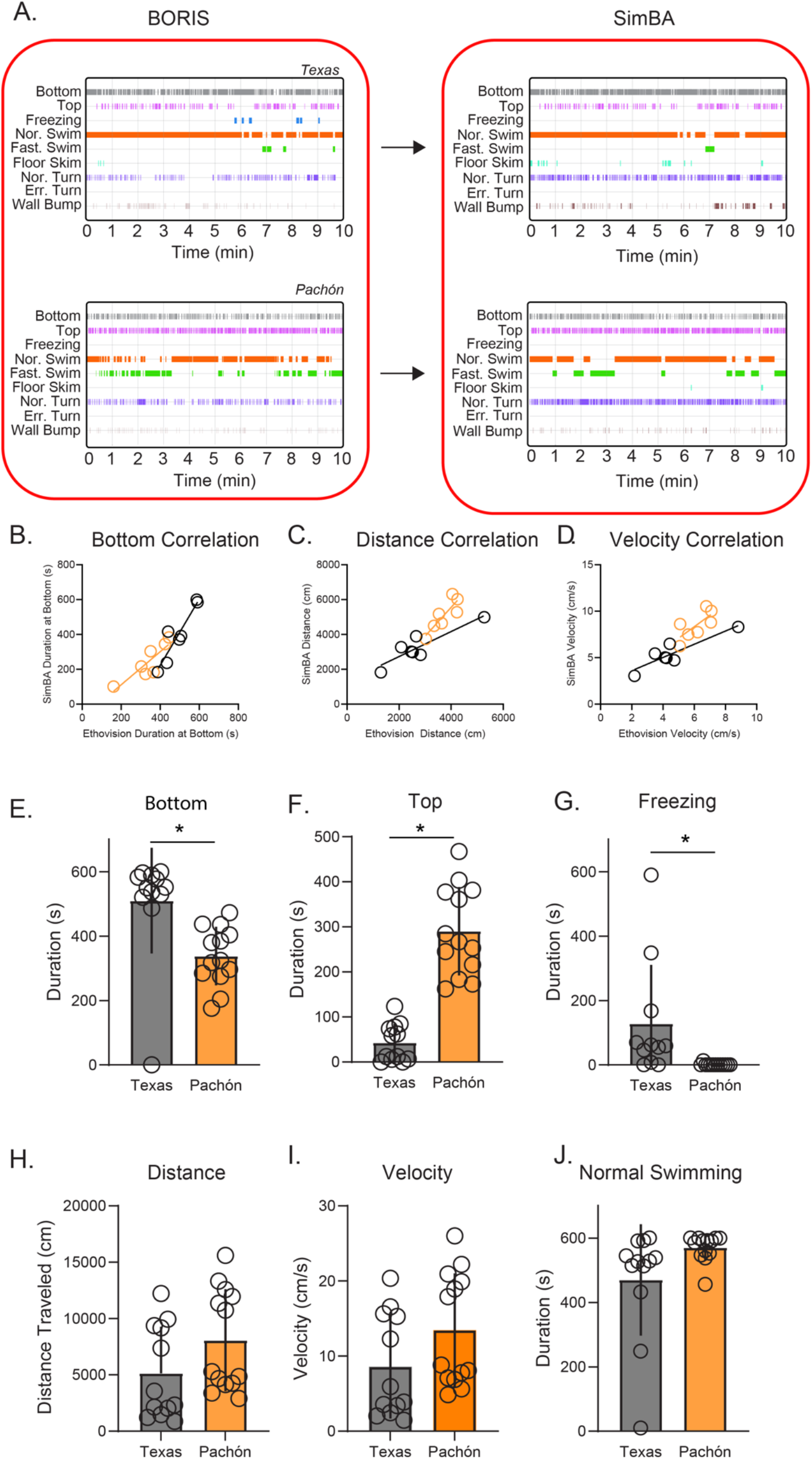
Behavioral classifiers suggest bottom-dwelling and freezing show differences between surface fish and cavefish. (A) Ethograms for surface fish (Texas) and cavefish (Pachón). A subset of the videos was annotated manually using BORIS. Manual annotations were fed into and trained with a behavioral classifier (SimBA). There was a concurrence between manual and automated predictions. (B-D) Results from Ethovision XT 13 and SimBA showed significant correlations for (B) time spent at the bottom (surface r^2^ = 0.87, p = 0.002; cave r^2^ = 0.75, p = 0.01), (C) distance traveled (surface r^2^ = 0.80, p = 0.006, cave r^2^ = 0.75, p = 0.01), and velocity (surface r^2^ = 0.80, p = 0.006, cave r^2^ = 0.53, p = 0.06). (E-F) Quantification of total duration spent in the (E) bottom or (F) top of the tank reveals significant differences between surface and cavefish (bottom: p = 0.003; top: p < 0.0001). (G) Quantification of freezing revealed that surface morphs significantly experience longer periods of immobility than cave morphs (p = 0.02). (H-I) Quantification of distance and velocity. They were found to be increased in the cave morph, but it did not yield significance (p = 0.1049; p = 0.1049). (J) Quantification of time spent normal swimming. It was observed that cave morphs spent more time normal swimming compared to surface (p = 0.054). Panel B-D compares data from Fig 1 with surface n=7 and Pachón n=7. Panels E-J contains Surface=12 and Pachón=13.

To confirm the reliability of SimBA, we compared results acquired with Ethovision to those produced by SimBA (Fig 3B-D). The same sets of videos were analyzed by both approaches and the results were compared. We observed strong correlations between values obtained by the two methods, confirming the accuracy of SimBA’s predictions (Fig. 3B-D; bottom-dwelling: surface r^2^ = 0.87, p = 0.002; cave r^2^ = 0.75, p = 0.01; distance traveled: surface r^2^ = 0.80, p = 0.006, cave r^2^ = 0.75, p = 0.01; average velocity: surface r^2^ = 0.80, p = 0.006, cave r^2^ = 0.53, p = 0.06).

Next, we analyzed differences in each of the nine behavioral parameters (Table 1) between surface and Pachón cave fish. Surface fish spent significantly more time in the bottom half of the tank and less time in the top half, consistent with previous findings (Fig. 3E, F). Freezing behavior occurred significantly more often in surface fish than in cavefish (Fig. 3G), supporting our previous observation that surface fish display more pronounced stress-associated behaviors. Other behavioral measures, such as total distance traveled, average velocity, and normal swimming, did not differ significantly between the two morphs (Fig. 3H-J, p>0.05). These findings reveal that some, but not all, behavioral parameters assessed were different between these two morphs, and highlight the need for detailed assessment of complex behaviors with multiple parameters.

### High throughput analysis of F2 hybrids reveals stress behaviors that co-evolved

*S*urface and cave morphs of *A. mexicanus* are interfertile providing the opportunity to examine the genetic relationship between distinct traits (Jeffery, 2001). By producing F2 surface x cave hybrids, phenotyping multiple traits, and assessing which traits correlate with one another, we can infer which traits are genetically linked. To investigate relationships among stress-related behaviors, we recorded the behavior of 50 F2 hybrid fish during the novel tank assay. Using our trained behavioral classifier, SimBA, we predicted the duration and frequency of specific behaviors for each fish. These predicted measures were then analyzed to calculate pairwise correlations between all behaviors, resulting in a pair-wise correlation matrix (Fig. 4A). Hierarchical clustering was subsequently performed on the correlation matrix to identify groups of behaviors that were highly correlated with one another (Fig. 4B). The clustering analysis revealed two distinct groups of traits (Fig 4B). Notably, duration of normal swimming was not significantly correlated with any other measured behavior and served as an outgroup. In contrast, freezing duration and bottom-dwelling duration were strongly correlated, forming a distinct cluster. This suggests that these two behaviors are closely associated and may reflect shared genetic underpinnings or a coordinated physiological response to stress. All other behaviors grouped into a separate cluster, indicating a different set of potentially co-regulated traits. These findings highlight freezing and bottom-dwelling as core components of an evolved stress-response phenotype, especially to low-predation environments. The strong correlation between these behaviors suggests they may arise from overlapping mechanisms, such as shared neural circuits or hormonal pathways, and could represent an integrated strategy for responding to stressors. Together, this analysis provides new insights into the organization and potential genetic basis of stress-related behaviors in *Astyanax mexicanus*

**Fig 4.**
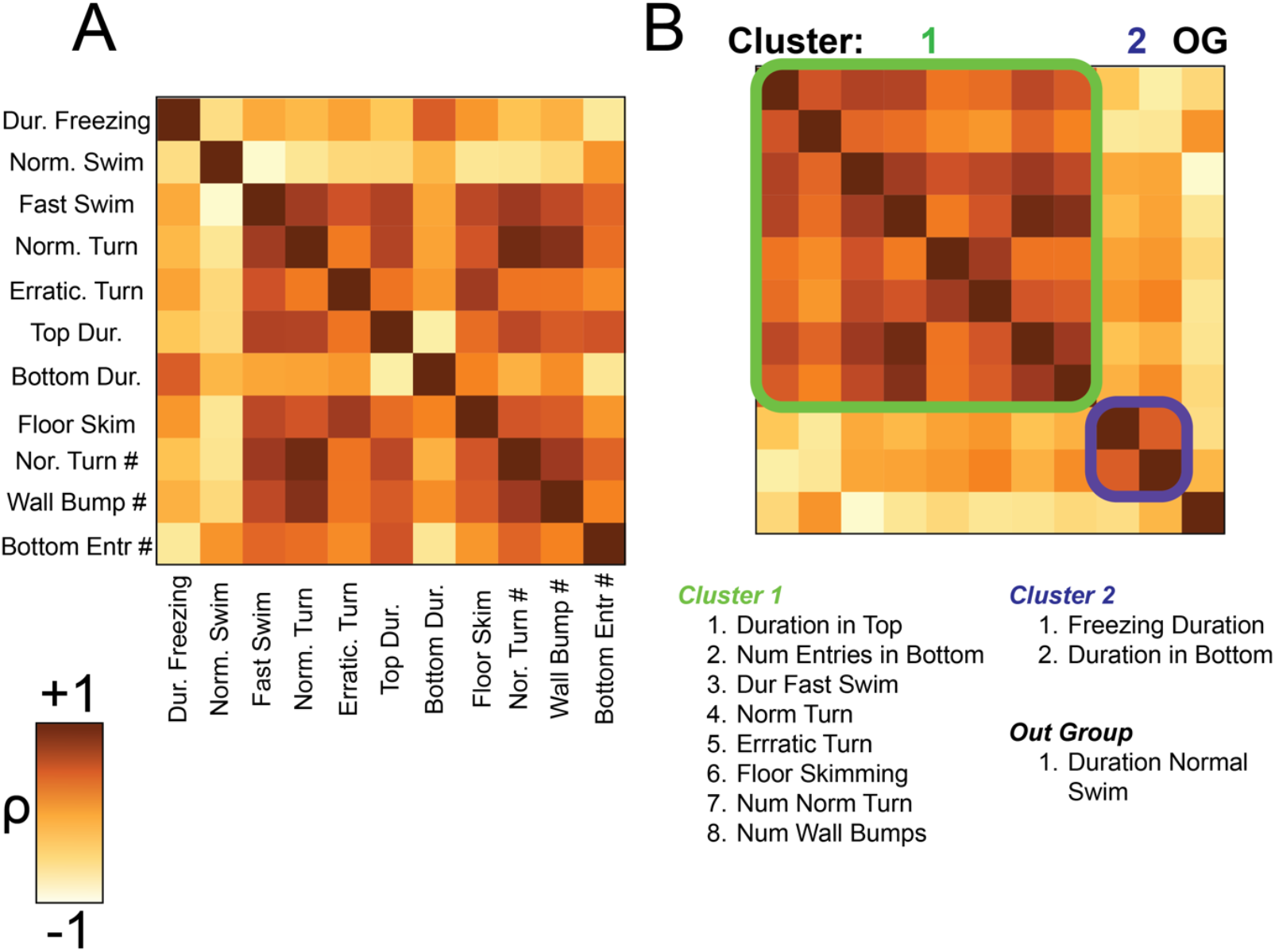
Stress behaviors surface x cave F2 hybrids appear to cluster into two main groups, suggesting that these traits co-segregate. (A) Heatmap displaying the pairwise correlations (ρ\rho) between behavioral traits. The intensity of the color represents the strength and direction of correlation. (B) Hierarchical clustering of the correlation matrix identified two primary clusters. Cluster 1 (green) comprises traits such as duration in the top, fast swimming, erratic turns, and wall bumping, indicative of exploratory or active behaviors. Cluster 2 (blue) includes freezing duration and time spent at the bottom, which are associated with stress-like behaviors. Duration normal swimming (OG, outgroup) showed no significant correlation with other traits. These results suggest co-segregation of stress-related behaviors in the F2 population. n=50

## Discussion

The current study investigated evolutionary differences in stress-related behaviors between surface-dwelling and cave-dwelling populations of *Astyanax mexicanus*. The primary goal was to understand how evolutionary adaptations influence stress responses by comparing these morphs in a novel tank test. Using pose-tracking and machine learning tools, we analyzed a range of stress-related behaviors, including freezing, bottom-dwelling, and hyperactivity. Our results confirmed that surface fish exhibit behaviors indicative of elevated stress, such as significantly increased time spent at the bottom of the tank and more frequent freezing. In contrast, cavefish demonstrated reduced stress-like behaviors, characterized by less freezing and greater exploration. Extending this analysis to F2 hybrids, we found that freezing and bottom-dwelling behaviors co-vary, suggesting these traits share genetic underpinnings or reflect related physiological processes.

### Stress Behaviors in an Evolutionary and Ecological Context

Stress responses are fundamental to survival, but their expression in the wild varies depending on ecological conditions (Mateo, 2007; Fischer et al., 2014; Heinen-Kay et al., 2016). Surface fish inhabit predator-rich environments and exhibit heightened stress responses, such as prolonged bottom-dwelling and freezing. In contrast, cavefish, which evolved in predator-free environments, display reduced stress-like behaviors characterized by increased exploration and minimal freezing (Mitchell et al., 1977; Chin et al., 2018; Chin et al., 2020). These contrasting phenotypes align with theories of adaptive evolution, where stress sensitivity is maintained under high predation pressure but relaxed in predator-free contexts.

The observed reduction in stress responses among cavefish likely reflects a broader shift in life-history strategy. Without the constant threat of predation, energy that would otherwise be allocated to acute stress responses can be redirected toward behaviors such as foraging efficiency, reproduction, or exploratory behavior. These findings are consistent with behavioral syndromes in other species, where boldness and reduced vigilance co-evolve in low-risk environments (Huntingford, 1976; Riechert and Hall, 2000; Dingemanse et al., 2004). For instance, similar patterns are observed in three-spined sticklebacks and desert spider populations, supporting the hypothesis that relaxed predation pressure universally selects for reduced stress responses (Huntingford, 1976; Riechert and Hall, 2000; Dingemanse et al., 2004).

The observed differences in stress-related behaviors also provide insight into the integration of these traits within broader behavioral syndromes. In surface fish, heightened stress responses may co-evolve with risk-averse behaviors to mitigate predation threats, while cavefish exhibit a bold, exploratory phenotype consistent with a predator-free habitat. These findings suggest that stress-related behaviors are not independent traits but part of a coordinated suite of ecological and evolutionary adaptations.

### Genetic Basis of Stress-Related Behaviors

The clustering of the stress-related traits freezing and bottom-dwelling in F2 hybrids suggests shared genetic underpinnings. This co-segregation may arise from pleiotropy, where a single gene influences multiple traits, or genetic linkage, where genes controlling these behaviors are inherited together. One possible mechanism is the role of cortisol as a common initiator of multiple stress-related responses, which could explain the co-occurrence of freezing and bottom-dwelling in hybrids. Elevated cortisol levels are known to induce freezing behavior and reduced exploration in zebrafish, and similar mechanisms may be at play in *Astyanax* mexicanus. These findings align with studies in zebrafish, where stress-related behaviors are mediated by glucocorticoid receptor signaling pathways (De Marco et al., 2013; Ziv et al., 2013; Chin et al., 2022). In surface fish, heightened stress responses likely involve increased glucocorticoid signaling, whereas cavefish may have evolved reduced sensitivity to cortisol as part of their adaptation to a predator-free environment. Prior work has shown that cavefish exhibit attenuated cortisol responses to stressors, suggesting modifications in glucocorticoid receptor pathways or feedback mechanisms (Gallo and Jeffery, 2012; Chin et al., 2018; Chin et al., 2020). This could underlie the reduced freezing and bottom-dwelling behaviors observed in cave populations. Together, these results suggest that the diminished stress phenotype in cavefish may reflect genetic and physiological shifts in glucocorticoid signaling, which aligns with relaxed selection pressures on stress-related traits in the absence of predators. Further genetic analyses will be necessary to determine whether specific modifications in glucocorticoid receptors or downstream signaling components contribute to these evolved differences in stress behavior.

### Computational Approaches for Behavioral Analysis

The application of tracking and machine learning tools, including pose-estimation and behavioral classifiers, has transformed the study behavior (Branson et al., 2009; Perez-Escudero et al., 2014; Mathis et al., 2018; Patch et al., 2022; Chen et al., 2023; Goodwin et al., 2024), but has not to date been applied to non-traditional evolutionary systems. By enabling precise, unbiased analysis of complex behaviors across large datasets, these methods overcome the limitations of manual scoring and provide a more comprehensive picture of stress phenotypes. Our application to *Astyanax* now permits the use of this established system to understand the evolution of complex behaviors. Behaviors such as freezing, bottom-dwelling, and hyperactivity, which often exhibit subtle differences, can now be quantified with greater accuracy and consistency. In the future, these approaches can be applied to myriad behaviors that differ between surface and cavefish, such as aggression and schooling/shoaling or other collective behaviors, allowing for an unprecedented look into how these complex traits evolve (Patch et al., 2022; Rodriguez-Morales et al., 2022; Paz et al., 2023). Additionally, they can be applied to diverse groups of animals and emerging non-traditional models. Moreover, by coupling automated behavioral phenotyping with genetic analyses, researchers can identify quantitative trait loci (QTL) associated with stress responses and explore the molecular pathways underlying these traits. Such approaches open new avenues for investigating the genetic basis of behavioral adaptation in natural systems and for extending these methods to other species with ecological contrasts.

## Conclusion

Our findings provide compelling evidence that stress-related behaviors in *A. mexicanus* are shaped by ecological and evolutionary pressures, with surface fish displaying heightened stress responses in predator-rich environments and cavefish exhibiting reduced stress behaviors in predator-free habitats. By leveraging advanced computational tools, this study establishes a foundation for high-throughput analysis of behavioral traits and offers new opportunities to investigate the genetic architecture of stress responses in natural systems.

## Supporting information

SimBA Feature Extraction Script for Fish

## Acknowledgements

The authors would like to thank members of the Duboue lab for their help and support. Experiments were carried out by NP, RA, SC, and SM. NP and SRON performed the computational work in SimBA. NP analyzed all data. NP and ERD wrote the manuscript with feedback from all authors. All authors contributed to the design of the experiments, and planning and oversight of the work. This work was funded by grants from the NIH (R15-MH132057) and the BSF (2019262) to ERD. Some of this work was also funded by grants from NIH (R15-MH118625) to ERD and an NSF EDGE grant (1923372) to ERD, JEK and ACK.

