## Supplementary material for "Automated behavioral profiling using neural networks reveals differences in stress-like behavior between cave and surface-dwelling *Astyanax mexicanus*": SimBA Feature Extraction Script for Fish: fish_feature_extractor_2023_version_5.py

```
from __future__ import division

import glob
import math
import os
from itertools import combinations

import numpy as np
import pandas as pd
from joblib import Parallel, delayed
from numba import jit, prange
from scipy import stats
from scipy.signal import find_peaks
from scipy.spatial import ConvexHull
from scipy.spatial.qhull import QhullError
from scipy.stats import zscore
from statsmodels.stats.diagnostic import lilliefors

from simba.mixins.config_reader import ConfigReader
from simba.mixins.feature_extraction_mixin import FeatureExtractionMixin
from simba.utils.enums import Paths
from simba.utils.printing import SimbaTimer
from simba.utils.read_write import (check_if_filepath_list_is_empty,
                                    get_fn_ext, read_config_file, read_df,
                                    read_project_path_and_file_type,
                                    read_video_info, write_df)

TAIL_BP_NAMES = ["objectA", "peduncle_base"]
CENTER_BP_NAMES = ["midpoint"]
MOUTH = ["mouth"]

ANGULAR_DISPERSION_S = [10, 5, 2, 1, 0.5, 0.25]


class FishFeatureExtractor(ConfigReader, FeatureExtractionMixin):
    def __init__(self, config_path: str):
        ConfigReader.__init__(self, config_path=config_path)
        FeatureExtractionMixin.__init__(self, config_path=config_path)
        self.timer = SimbaTimer()
        self.timer.start_timer()
        self.compass_brackets = ["N", "NE", "E", "SE", "S", "SW", "W", "NW", "N"]
        self.compass_brackets_long = [
            "Direction_N",
            "Direction_NE",
            "Direction_E",
            "Direction_SE",
            "Direction_S",
            "Direction_SW",
            "Direction_W",
            "Direction_NW",
        ]
        self.compass_brackets_digits = ["0", "1", "2", "3", "4", "5", "6", "7", "0"]
        self.config = read_config_file(config_path=config_path)
        self.project_path, self.file_type = read_project_path_and_file_type(
            config=self.config
        )
        self.input_file_dir = os.path.join(
            self.project_path, Paths.OUTLIER_CORRECTED.value
        )
        self.save_dir = os.path.join(
            self.project_path, Paths.FEATURES_EXTRACTED_DIR.value
        )
        self.video_info_path = os.path.join(self.project_path, Paths.VIDEO_INFO.value)
        self.video_info_df = pd.read_csv(self.video_info_path)
        bp_names_path = os.path.join(self.project_path, Paths.BP_NAMES.value)
        self.bp_names = list(pd.read_csv(bp_names_path, header=None)[0])
        self.col_headers_shifted = []
        for bp in self.bp_names:
            self.col_headers_shifted.extend(
                (bp + "_x_shifted", bp + "_y_shifted", bp + "_p_shifted")
            )
        self.x_y_cols = []
        self.x_cols_shifted, self.y_cols_shifted = [], []
        for x_name, y_name in zip(self.x_cols, self.y_cols):
            self.x_y_cols.extend((x_name, y_name))
            self.x_cols_shifted.append(x_name + "_shifted")
            self.y_cols_shifted.append(y_name + "_shifted")

        self.roll_windows_values = [
            75,
            50,
            25,
            20,
            15,
            10,
            4,
            2,
        ]
        self.files_found = glob.glob(
            self.input_file_dir + "/*.{}".format(self.file_type)
        )
        check_if_filepath_list_is_empty(
            filepaths=self.files_found,
            error_msg="SIMBA ERROR: No file in {} directory".format(
                self.input_file_dir
            ),
        )
        print(
            "Extracting features from {} {}...".format(
                str(len(self.files_found)), "file(s)"
            )
        )

        for file_path in self.files_found:
            video_timer = SimbaTimer(start=True)
            video_timer.start_timer()
            dir_name, file_name, ext = get_fn_ext(file_path)
            self.save_path = os.path.join(self.save_dir, os.path.basename(file_path))
            video_info, self.px_per_mm, self.fps = read_video_info(
                self.video_info_df, file_name
            )
            self.video_width, self.video_height = (
                video_info["Resolution_width"].values,
                video_info["Resolution_height"].values,
            )
            self.angular_dispersion_windows = []
            for i in range(len(ANGULAR_DISPERSION_S)):
                self.angular_dispersion_windows.append(
                    int(self.fps * ANGULAR_DISPERSION_S[i])
                )

            self.csv_df = (
                read_df(file_path, self.file_type).fillna(0).apply(pd.to_numeric)
            )
            try:
                self.csv_df.columns = self.bp_headers
            except ValueError:
                msg = f"ERROR: Data contains the following fields: {self.csv_df.columns}. \n SimBA wants to use the following field names {self.bp_header_list}"
                print(msg)
                raise ValueError(msg)

            csv_df_shifted = self.csv_df.shift(periods=1)
            csv_df_shifted.columns = self.col_headers_shifted
            self.csv_df_combined = pd.concat(
                [self.csv_df, csv_df_shifted], axis=1, join="inner"
            ).fillna(0)
            self.calc_X_relative_to_Y_movement()
            self.calc_movement()
            self.calc_X_relative_to_Y_movement_rolling_windows()
            self.calc_velocity()
            self.calc_acceleration()
            self.calc_rotation()
            self.calc_N_degree_direction_switches()
            self.bouts_in_same_direction()
            self.calc_45_degree_direction_switches()
            self.hot_end_encode_compass()
            self.calc_directional_switches_in_rolling_windows()
            self.calc_angular_dispersion()
            self.calc_border_distances()
            self.calc_distances_between_body_part()
            self.calc_convex_hulls()
            self.pose_confidence_probabilities()
            self.distribution_tests()
            self.save_file()
            video_timer.stop_timer()
            print(
                f"Features extracted for video {file_name} (elapsed time {video_timer.elapsed_time_str}s)..."
            )

        self.timer.stop_timer()
        print(
            f"Features extracted for all {str(len(self.files_found))} files, data saved in project_folder/csv/features_extracted directory (elapsed time {self.timer.elapsed_time_str}s)"
        )

    def angle2pt_degrees(self, ax, ay, bx, by):
        angle_degrees = math.degrees(math.atan2(ax - bx, by - ay))
        return angle_degrees + 360 if angle_degrees < 0 else angle_degrees

    def angle2pt_radians(self, degrees):
        angle_radians = degrees * math.pi / 180
        return angle_radians

    def angle2pt_sin(self, angle_radians):
        angle_sin = math.sin(angle_radians)
        return angle_sin

    def angle2pt_cos(self, angle_radians):
        angle_cos = math.cos(angle_radians)
        return angle_cos

    @staticmethod
    @jit(nopython=True)
    def count_values_in_range(data: np.array, ranges: np.array):
        results = np.full((data.shape[0], ranges.shape[0]), 0)
        for i in prange(data.shape[0]):
            for j in prange(ranges.shape[0]):
                lower_bound, upper_bound = ranges[j][0], ranges[j][1]
                results[i][j] = data[i][
                    np.logical_and(data[i] >= lower_bound, data[i] <= upper_bound)
                ].shape[0]
        return results

    @staticmethod
    def convex_hull_calculator_mp(arr: np.array, px_per_mm: float) -> float:
        arr = np.unique(arr, axis=0).astype(int)
        if arr.shape[0] < 3:
            return 0
        for i in range(1, arr.shape[0]):
            if (arr[i] != arr[0]).all():
                try:
                    return ConvexHull(arr, qhull_options="En").area / px_per_mm
                except QhullError:
                    return 0
            else:
                pass
        return 0

    @staticmethod
    @jit(nopython=True)
    def euclidian_distance_calc(bp1xVals, bp1yVals, bp2xVals, bp2yVals):
        return np.sqrt((bp1xVals - bp2xVals) ** 2 + (bp1yVals - bp2yVals) ** 2)

    @staticmethod
    @jit(nopython=True)
    def angular_dispersion(cumsum_cos_np, cumsum_sin_np):
        out_array = np.empty((cumsum_cos_np.shape))
        for index in range(cumsum_cos_np.shape[0]):
            X, Y = cumsum_cos_np[index] / (index + 1), cumsum_sin_np[index] / (
                index + 1
            )
            out_array[index] = math.sqrt(X**2 + Y**2)
        return out_array

    def windowed_frequentist_distribution_tests(
        self, data: np.array, feature_name: str, fps: int
    ):
        """
        Helper to compare feature value distributions in 1s sequential time-bins: Kolmogorov-Smirnov and T-tests
        Compares the feature values against a normal distribution: Lillefors, Shapiro.
        Find the number of peaks in *rolling* 1s long feature window.
        """

        (ks_results,) = (np.full((data.shape[0]), -1.0),)
        t_test_results = np.full((data.shape[0]), -1.0)
        lillefors_results = np.full((data.shape[0]), -1.0)
        shapiro_results = np.full((data.shape[0]), -1.0)
        peak_cnt_results = np.full((data.shape[0]), -1.0)

        for i in range(fps, data.shape[0] - fps, fps):
            bin_1_idx, bin_2_idx = [i - fps, i], [i, i + fps]
            bin_1_data, bin_2_data = (
                data[bin_1_idx[0] : bin_1_idx[1]],
                data[bin_2_idx[0] : bin_2_idx[1]],
            )
            ks_results[i : i + fps + 1] = stats.ks_2samp(
                data1=bin_1_data, data2=bin_2_data
            ).statistic
            t_test_results[i : i + fps + 1] = stats.ttest_ind(
                bin_1_data, bin_2_data
            ).statistic

        for i in range(0, data.shape[0] - fps, fps):
            lillefors_results[i : i + fps + 1] = lilliefors(data[i : i + fps])[0]
            shapiro_results[i : i + fps + 1] = stats.shapiro(data[i : i + fps])[0]

        rolling_idx = np.arange(fps)[None, :] + 1 * np.arange(data.shape[0])[:, None]
        for i in range(rolling_idx.shape[0]):
            bin_start_idx, bin_end_idx = rolling_idx[i][0], rolling_idx[i][-1]
            peaks, _ = find_peaks(data[bin_start_idx:bin_end_idx], height=0)
            peak_cnt_results[i] = len(peaks)

        columns = [
            f"{feature_name}_KS",
            f"{feature_name}_TTEST",
            f"{feature_name}_LILLEFORS",
            f"{feature_name}_SHAPIRO",
            f"{feature_name}_PEAK_CNT",
        ]
        return pd.DataFrame(
            np.column_stack(
                (
                    ks_results,
                    t_test_results,
                    lillefors_results,
                    shapiro_results,
                    peak_cnt_results,
                )
            ),
            columns=columns,
        ).round(4)

    @staticmethod
    @jit(nopython=True)
    def consecutive_frames_in_same_compass_direction(direction: np.array):
        results = np.full((direction.shape[0], 1), -1)
        cnt, results[0], last_direction = 0, 0, direction[0]
        for i in prange(1, direction.shape[0]):
            if direction[i] == last_direction:
                cnt += 1
            else:
                cnt = 0
            results[i] = cnt
            last_direction = direction[i]
        return results.flatten()

    @staticmethod
    @jit(nopython=True, fastmath=True)
    def framewise_degree_shift(clockwise_angle: np.array):
        degree_shift = np.full((clockwise_angle.shape[0], 1), np.nan)
        cnt, degree_shift[0], last_angle = 0, 0, clockwise_angle[0]
        for i in prange(1, clockwise_angle.shape[0]):
            degree_shift[i] = math.atan2(
                math.sin(clockwise_angle[i] - last_angle),
                math.cos(clockwise_angle[i] - last_angle),
            )
        return np.absolute(degree_shift.flatten())

    def bouts_in_same_direction(self):
        self.csv_df_combined["Consecutive_ms_in_same_compass_direction"] = (
            self.consecutive_frames_in_same_compass_direction(
                direction=self.csv_df_combined["Compass_digit"].values.astype(int)
            )
            / self.fps
        )
        self.csv_df_combined["Consecutive_ms_in_same_compass_direction_zscore"] = (
            zscore(
                self.csv_df_combined["Consecutive_ms_in_same_compass_direction"].values
            )
        )
        for window in self.roll_windows_values:
            self.csv_df_combined[f"Unique_compass_directions_in_{window}_window"] = (
                self.csv_df_combined["Compass_digit"]
                .astype(int)
                .rolling(window, min_periods=1)
                .apply(lambda x: len(np.unique(x)))
                .astype(int)
            )
        framewise_degree_shift = pd.Series(
            self.framewise_degree_shift(
                clockwise_angle=self.csv_df_combined["Clockwise_angle_degrees"].values
            )
        )
        for window in self.roll_windows_values:
            self.csv_df_combined[f"Degree_shift_{window}_mean"] = (
                framewise_degree_shift.rolling(window, min_periods=1).mean()
            )
            self.csv_df_combined[f"Degree_shift_{window}_median"] = (
                framewise_degree_shift.rolling(window, min_periods=1).median()
            )
            self.csv_df_combined[f"Degree_shift_{window}_sum"] = (
                framewise_degree_shift.rolling(window, min_periods=1).sum()
            )
            self.csv_df_combined[f"Degree_shift_{window}_std"] = (
                framewise_degree_shift.rolling(window, min_periods=1).std()
            )

    def calc_angular_dispersion(self):
        dispersion_array = self.angular_dispersion(
            self.csv_df_combined["Angle_cos_cumsum"].values,
            self.csv_df_combined["Angle_sin_cumsum"].values,
        )
        self.csv_df_combined["Angular_dispersion"] = dispersion_array

        for win in range(len(self.angular_dispersion_windows)):
            col_name = "Angular_dispersion_window_" + str(
                self.angular_dispersion_windows[win]
            )
            self.csv_df_combined[col_name] = (
                self.csv_df_combined["Angular_dispersion"]
                .rolling(self.angular_dispersion_windows[win], min_periods=1)
                .mean()
            )

    def calc_X_relative_to_Y_movement(self):
        temp_df = pd.DataFrame()
        for bp in range(len(self.x_cols)):
            curr_x_col, curr_x_shifted_col, curr_y_col, curr_y_shifted_col = (
                self.x_cols[bp],
                self.x_cols_shifted[bp],
                self.y_cols[bp],
                self.y_cols_shifted[bp],
            )
            temp_df["x"] = (
                self.csv_df_combined[curr_x_col]
                - self.csv_df_combined[curr_x_shifted_col]
            )
            temp_df["y"] = (
                self.csv_df_combined[curr_y_col]
                - self.csv_df_combined[curr_y_shifted_col]
            )
            temp_df["Movement_{}_X_relative_2_Y".format(bp)] = (
                temp_df["x"] - temp_df["y"]
            )
            temp_df.drop(["x", "y"], axis=1, inplace=True)
        self.csv_df_combined["Movement_X_axis_relative_to_Y_axis"] = temp_df.sum(axis=1)

    def calc_movement(self):
        movement_cols = []
        for bp in self.bp_names:
            self.csv_df_combined[f"{bp}_movement"] = (
                self.euclidian_distance_calc(
                    self.csv_df_combined[f"{bp}_x"].values,
                    self.csv_df_combined[f"{bp}_y"].values,
                    self.csv_df_combined[f"{bp}_x_shifted"].values,
                    self.csv_df_combined[f"{bp}_y_shifted"].values,
                )
                / self.px_per_mm
            )
            movement_cols.append(f"{bp}_movement")
        self.csv_df_combined["Summed_movement"] = self.csv_df_combined[
            movement_cols
        ].sum(axis=1)

        for bp in self.bp_names:
            for window in self.roll_windows_values:
                self.csv_df_combined[f"{bp}_movement_{window}_mean"] = (
                    self.csv_df_combined[f"{bp}_movement"]
                    .rolling(window, min_periods=1)
                    .mean()
                )
                self.csv_df_combined[f"{bp}_movement_{window}_sum"] = (
                    self.csv_df_combined[f"{bp}_movement"]
                    .rolling(window, min_periods=1)
                    .sum()
                )

    def calc_X_relative_to_Y_movement_rolling_windows(self):
        for i in self.roll_windows_values:
            currentColName = f"Movement_X_axis_relative_to_Y_axis_mean_{i}"
            self.csv_df_combined[currentColName] = (
                self.csv_df_combined["Movement_X_axis_relative_to_Y_axis"]
                .rolling(i, min_periods=1)
                .mean()
            )
            currentColName = f"Movement_X_axis_relative_to_Y_axis_sum_{i}"
            self.csv_df_combined[currentColName] = (
                self.csv_df_combined["Movement_X_axis_relative_to_Y_axis"]
                .rolling(i, min_periods=1)
                .sum()
            )

    def calc_directional_switches_in_rolling_windows(self):
        for i in self.roll_windows_values:
            currentColName = f"Number_of_direction_switches_{i}"
            self.csv_df_combined[currentColName] = (
                self.csv_df_combined["Direction_switch"].rolling(i, min_periods=1).sum()
            )
            currentColName = f"Directionality_of_switches_switches_{i}"
            self.csv_df_combined[currentColName] = (
                self.csv_df_combined["Switch_direction_value"]
                .rolling(i, min_periods=1)
                .sum()
            )

    def calc_velocity(self):
        self.velocity_fields = []
        for bp in self.bp_names:
            self.csv_df_combined[f"{bp}_velocity"] = (
                self.csv_df_combined[bp + "_movement"]
                .rolling(int(self.fps), min_periods=1)
                .sum()
            )
            self.velocity_fields.append(bp + "_velocity")
        self.csv_df_combined["Bp_velocity_mean"] = self.csv_df_combined[
            self.velocity_fields
        ].mean(axis=1)
        self.csv_df_combined["Bp_velocity_stdev"] = self.csv_df_combined[
            self.velocity_fields
        ].std(axis=1)
        for i in self.roll_windows_values:
            self.csv_df_combined[f"Minimum_avg_bp_velocity_{i}_window"] = (
                self.csv_df_combined["Bp_velocity_mean"].rolling(i, min_periods=1).min()
            )
            self.csv_df_combined[f"Max_avg_bp_velocity_{i}_window"] = (
                self.csv_df_combined["Bp_velocity_mean"].rolling(i, min_periods=1).max()
            )
            self.csv_df_combined[
                f"Absolute_diff_min_max_avg_bp_velocity_{i}_window"
            ] = abs(
                self.csv_df_combined[f"Minimum_avg_bp_velocity_{i}_window"]
                - self.csv_df_combined[f"Max_avg_bp_velocity_{i}_window"]
            )

    def calc_acceleration(self):
        for i in self.roll_windows_values:
            acceleration_fields = []
            for bp in self.bp_names:
                self.csv_df_combined[f"{bp}_velocity_shifted"] = (
                    self.csv_df_combined[f"{bp}_velocity"]
                    .shift(i)
                    .fillna(self.csv_df_combined[f"{bp}_velocity"])
                )
                self.csv_df_combined[f"{bp}_acceleration_{i}_window"] = (
                    self.csv_df_combined[f"{bp}_velocity"]
                    - self.csv_df_combined[f"{bp}_velocity_shifted"]
                )
                self.csv_df_combined = self.csv_df_combined.drop(
                    [f"{bp}_velocity_shifted"], axis=1
                )
                acceleration_fields.append(f"{bp}_acceleration_{i}_window")
            self.csv_df_combined[f"Bp_acceleration_mean_{i}_window"] = (
                self.csv_df_combined[acceleration_fields].mean(axis=1)
            )
            self.csv_df_combined[f"Bp_acceleration_stdev_{i}_window"] = (
                self.csv_df_combined[acceleration_fields].std(axis=1)
            )
        for i in self.roll_windows_values:
            self.csv_df_combined[f"Min_avg_bp_acceleration_{i}_window"] = (
                self.csv_df_combined[f"Bp_acceleration_mean_{i}_window"]
                .rolling(i, min_periods=1)
                .mean()
            )
            self.csv_df_combined[f"Max_avg_bp_acceleration_{i}_window"] = (
                self.csv_df_combined[f"Bp_acceleration_mean_{i}_window"]
                .rolling(i, min_periods=1)
                .mean()
            )
            self.csv_df_combined[
                f"Absolute_diff_min_max_avg_bp_velocity_{i}_window"
            ] = abs(
                self.csv_df_combined[f"Min_avg_bp_acceleration_{i}_window"]
                - self.csv_df_combined[f"Max_avg_bp_acceleration_{i}_window"]
            )

    def calc_N_degree_direction_switches(self):
        degree_lk_180 = {"N": ["S"], "NE": ["SW"], "E": ["W"], "SE": ["NW"]}
        degree_lk_90 = {
            "N": ["W", "E"],
            "NE": ["NW", "SE"],
            "NW": ["SW", "NE"],
            "SW": ["NW", "SE"],
            "SE": ["NE", "SW"],
            "S": ["W", "E"],
            "E": ["N", "S"],
            "W": ["N", "S"],
        }
        dg_df = pd.DataFrame(self.csv_df_combined["Compass_direction"])
        for window in self.roll_windows_values:
            dg_df[f"Compass_direction_{window}"] = dg_df["Compass_direction"].shift(
                window
            )
            dg_df[f"Compass_direction_{window}"].fillna(
                dg_df["Compass_direction"], inplace=True
            )
            dg_df[f"180_degree_switch_{window}"] = 0
            dg_df[f"90_degree_switch_{window}"] = 0
            for k, v in degree_lk_180.items():
                for value in v:
                    dg_df.loc[
                        (dg_df["Compass_direction"] == k)
                        & (dg_df[f"Compass_direction_{window}"] == value),
                        f"180_degree_switch_{window}",
                    ] = 1
                    dg_df.loc[
                        (dg_df[f"Compass_direction_{window}"] == k)
                        & (dg_df["Compass_direction"] == value),
                        f"180_degree_switch_{window}",
                    ] = 1
            for k, v in degree_lk_90.items():
                for value in v:
                    dg_df.loc[
                        (dg_df["Compass_direction"] == k)
                        & (dg_df[f"Compass_direction_{window}"] == value),
                        f"90_degree_switch_{window}",
                    ] = 1
                    dg_df.loc[
                        (dg_df[f"Compass_direction_{window}"] == k)
                        & (dg_df["Compass_direction"] == value),
                        f"90_degree_switch_{window}",
                    ] = 1
            self.csv_df_combined[f"180_degree_switch_{window}"] = dg_df[
                f"180_degree_switch_{window}"
            ]
            self.csv_df_combined[f"90_degree_switch_{window}"] = dg_df[
                f"90_degree_switch_{window}"
            ]

    def calc_rotation(self):
        self.csv_df_combined["Clockwise_angle_degrees"] = self.csv_df_combined.apply(
            lambda x: self.angle2pt_degrees(
                x[CENTER_BP_NAMES[0] + "_x"],
                x[CENTER_BP_NAMES[0] + "_y"],
                x[TAIL_BP_NAMES[0] + "_x"],
                x[TAIL_BP_NAMES[0] + "_y"],
            ),
            axis=1,
        )
        self.csv_df_combined["Angle_radians"] = self.angle2pt_radians(
            self.csv_df_combined["Clockwise_angle_degrees"]
        )
        self.csv_df_combined["Angle_sin"] = self.csv_df_combined.apply(
            lambda x: self.angle2pt_sin(x["Angle_radians"]), axis=1
        )
        self.csv_df_combined["Angle_cos"] = self.csv_df_combined.apply(
            lambda x: self.angle2pt_cos(x["Angle_radians"]), axis=1
        )
        self.csv_df_combined["Angle_sin_cumsum"] = self.csv_df_combined[
            "Angle_sin"
        ].cumsum()
        self.csv_df_combined["Angle_cos_cumsum"] = self.csv_df_combined[
            "Angle_cos"
        ].cumsum()
        compass_lookup = list(
            round(self.csv_df_combined["Clockwise_angle_degrees"] / 45)
        )
        compass_lookup = [int(i) for i in compass_lookup]
        compasFaceList_bracket, compasFaceList_digit = [], []
        for compasDirection in compass_lookup:
            compasFaceList_bracket.append(self.compass_brackets[compasDirection])
            compasFaceList_digit.append(self.compass_brackets_digits[compasDirection])
        self.csv_df_combined["Compass_direction"] = compasFaceList_bracket
        self.csv_df_combined["Compass_digit"] = compasFaceList_digit
        for i in self.roll_windows_values:
            column_name = f"Mean_angle_time_window_{i}"
            self.csv_df_combined[column_name] = (
                self.csv_df_combined["Clockwise_angle_degrees"]
                .rolling(i, min_periods=1)
                .mean()
            )

    def hot_end_encode_compass(self):
        compass_hot_end = pd.get_dummies(
            self.csv_df_combined["Compass_direction"], prefix="Direction"
        )
        compass_hot_end = compass_hot_end.T.reindex(
            self.compass_brackets_long
        ).T.fillna(0)
        self.csv_df_combined = pd.concat(
            [self.csv_df_combined, compass_hot_end], axis=1
        )

    def calc_45_degree_direction_switches(self):
        self.grouped_df = pd.DataFrame()
        v = (
            self.csv_df_combined["Compass_digit"]
            != self.csv_df_combined["Compass_digit"].shift()
        ).cumsum()
        u = self.csv_df_combined.groupby(v)["Compass_digit"].agg(["all", "count"])
        m = u["all"] & u["count"].ge(1)
        self.grouped_df["groups"] = self.csv_df_combined.groupby(v).apply(
            lambda x: (x.index[0], x.index[-1])
        )[m]
        currdirectionList, DirectionSwitchIndexList, currdirectionListValue = [], [], []
        for indexes, row in self.grouped_df.iterrows():
            currdirectionList.append(
                self.csv_df_combined.loc[row["groups"][0]]["Compass_direction"]
            )
            DirectionSwitchIndexList.append(row["groups"][1])
            currdirectionListValue.append(
                self.csv_df_combined.loc[row["groups"][0]]["Compass_digit"]
            )
        self.grouped_df["Direction_switch"] = currdirectionList
        self.grouped_df["Direction_value"] = currdirectionListValue
        self.csv_df_combined.loc[DirectionSwitchIndexList, "Direction_switch"] = 1
        self.csv_df_combined["Compass_digit_shifted"] = self.csv_df_combined[
            "Compass_digit"
        ].shift(-1)
        self.csv_df_combined = self.csv_df_combined.fillna(0)
        self.csv_df_combined["Switch_direction_value"] = self.csv_df_combined.apply(
            lambda x: self.calc_switch_direction(
                x["Compass_digit_shifted"], x["Compass_digit"]
            ),
            axis=1,
        )

    def calc_switch_direction(self, compass_digit_shifted, compass_digit):
        if (compass_digit_shifted == "0") and (compass_digit == "7"):
            return 1
        else:
            return int(compass_digit_shifted) - int(compass_digit)

    def calc_border_distances(self):
        for bp in self.bp_names:
            self.csv_df_combined[f"{bp}_distance_to_left_border"] = (
                self.csv_df_combined[f"{bp}_x"] / self.px_per_mm
            )
            self.csv_df_combined[f"{bp}_distance_to_right_border"] = (
                self.video_width - self.csv_df_combined[f"{bp}_x"]
            ) / self.px_per_mm
            self.csv_df_combined[f"{bp}_distance_to_top_border"] = (
                self.csv_df_combined[f"{bp}_y"] / self.px_per_mm
            )
            self.csv_df_combined[f"{bp}_distance_to_bottom_border"] = (
                self.video_height - self.csv_df_combined[f"{bp}_y"]
            ) / self.px_per_mm

        for side in ["left", "right", "top", "bottom"]:
            side_col_names = [
                c
                for c in self.csv_df_combined.columns
                if f"distance_to_{side}_border" in c
            ]
            self.csv_df_combined[f"Mean_bp_distance_to_{side}_border"] = (
                self.csv_df_combined[side_col_names].mean(axis=1)
            )
            for window in self.roll_windows_values:
                self.csv_df_combined[f"Mean_bp_distance_to_{side}_border_{window}"] = (
                    self.csv_df_combined[f"Mean_bp_distance_to_{side}_border"]
                    .rolling(window, min_periods=1)
                    .mean()
                )
                self.csv_df_combined[f"Std_bp_distance_to_{side}_border_{window}"] = (
                    self.csv_df_combined[f"Mean_bp_distance_to_{side}_border"]
                    .rolling(window, min_periods=1)
                    .std()
                )
                try:
                    # self.csv_df_combined[f'Kurtosis_bp_distance_to_{side}_border_{window}'] = self.csv_df_combined[f'Mean_bp_distance_to_{side}_border'].rolling(window, min_periods=window).kurt()
                    self.csv_df_combined[
                        f"Skew_bp_distance_to_{side}_border_{window}"
                    ] = (
                        self.csv_df_combined[f"Mean_bp_distance_to_{side}_border"]
                        .rolling(window, min_periods=window)
                        .skew()
                    )
                except:
                    # self.csv_df_combined[f'Kurtosis_bp_distance_to_{side}_border_{window}'] = -1
                    self.csv_df_combined[
                        f"Skew_bp_distance_to_{side}_border_{window}"
                    ] = -1

    def calc_distances_between_body_part(self):
        two_point_combs = np.array(list(combinations(self.bp_names, 2)))
        distance_fields = []
        for bps in two_point_combs:
            self.csv_df_combined[f"Distance_{bps[0]}_{bps[1]}"] = (
                self.euclidian_distance_calc(
                    self.csv_df_combined[bps[0] + "_x"].values,
                    self.csv_df_combined[bps[0] + "_y"].values,
                    self.csv_df_combined[bps[1] + "_x"].values,
                    self.csv_df_combined[bps[1] + "_y"].values,
                )
                / self.px_per_mm
            )
            distance_fields.append(f"Distance_{bps[0]}_{bps[1]}")

        for distance_field in distance_fields:
            for window in self.roll_windows_values:
                self.csv_df_combined[f"{distance_field}_mean_{window}"] = (
                    self.csv_df_combined[distance_field]
                    .rolling(window, min_periods=1)
                    .mean()
                )
                self.csv_df_combined[f"{distance_field}_std_{window}"] = (
                    self.csv_df_combined[distance_field]
                    .rolling(window, min_periods=1)
                    .std()
                )
                try:
                    self.csv_df_combined[f"{distance_field}_skew_{window}"] = (
                        self.csv_df_combined[distance_field]
                        .rolling(window, min_periods=1)
                        .skew()
                    )
                    self.csv_df_combined[f"{distance_field}_kurtosis_{window}"] = (
                        self.csv_df_combined[distance_field]
                        .rolling(window, min_periods=1)
                        .kurt()
                    )
                except:
                    self.csv_df_combined[f"{distance_field}_skew_{window}"] = -1
                    self.csv_df_combined[f"{distance_field}_kurtosis_{window}"] = -1

    def calc_convex_hulls(self):
        fish_array = np.reshape(
            self.csv_df[self.x_y_cols].values, (len(self.csv_df / 2), -1, 2)
        )
        self.csv_df_combined["Convex_hull"] = Parallel(
            n_jobs=-1, verbose=0, backend="threading"
        )(
            delayed(self.convex_hull_calculator_mp)(x, self.px_per_mm)
            for x in fish_array
        )
        for window in self.roll_windows_values:
            self.csv_df_combined[f"Convex_hull_mean_{window}_window"] = (
                self.csv_df_combined["Convex_hull"]
                .rolling(window, min_periods=1)
                .mean()
            )
            self.csv_df_combined[f"Convex_hull_std_{window}_window"] = (
                self.csv_df_combined["Convex_hull"].rolling(window, min_periods=1).std()
            )
            self.csv_df_combined[f"Convex_hull_min_{window}_window"] = (
                self.csv_df_combined["Convex_hull"].rolling(window, min_periods=1).min()
            )
            self.csv_df_combined[f"Convex_hull_max_{window}_window"] = (
                self.csv_df_combined["Convex_hull"].rolling(window, min_periods=1).max()
            )
            self.csv_df_combined[
                f"Absolute_diff_min_max_convex_hull_{window}_window"
            ] = abs(
                self.csv_df_combined[f"Convex_hull_min_{window}_window"]
                - self.csv_df_combined[f"Convex_hull_max_{window}_window"]
            )
            try:
                self.csv_df_combined[f"Convex_hull_skew_{window}"] = (
                    self.csv_df_combined["Convex_hull"]
                    .rolling(window, min_periods=1)
                    .skew()
                )
                self.csv_df_combined[f"Convex_hull_kurtosis_{window}"] = (
                    self.csv_df_combined["Convex_hull"]
                    .rolling(window, min_periods=1)
                    .kurt()
                )
            except:
                self.csv_df_combined[f"Convex_hull_skew_{window}"] = -1
                self.csv_df_combined[f"Convex_hull_kurtosis_{window}"] = -1

    def distribution_tests(self):
        distribution_features = [
            "Mean_bp_distance_to_left_border",
            "Mean_bp_distance_to_right_border",
            "Mean_bp_distance_to_top_border",
            "Mean_bp_distance_to_bottom_border",
            "Bp_velocity_mean",
            "Bp_acceleration_mean_25_window",
            "Clockwise_angle_degrees",
            "Convex_hull",
            "Sum_probabilities",
            "Consecutive_ms_in_same_compass_direction",
        ]

        for feature_name in distribution_features:
            results = self.windowed_frequentist_distribution_tests(
                data=self.csv_df_combined[feature_name].values,
                feature_name=feature_name,
                fps=int(self.fps),
            )
            self.csv_df_combined = pd.concat([self.csv_df_combined, results], axis=1)

    def pose_confidence_probabilities(self):
        self.csv_df_combined["Sum_probabilities"] = self.csv_df_combined[
            self.p_cols
        ].sum(axis=1)
        self.csv_df_combined["Sum_probabilities_deviation"] = (
            self.csv_df_combined["Sum_probabilities"].mean()
            - self.csv_df_combined["Sum_probabilities"]
        )
        p_brackets_results = pd.DataFrame(
            self.count_values_in_range(
                data=self.csv_df_combined.filter(self.p_cols).values,
                ranges=np.array(
                    [
                        [0.0, 0.1],
                        [0.000000000, 0.5],
                        [0.000000000, 0.75],
                        [0.000000000, 0.95],
                        [0.000000000, 0.99],
                    ]
                ),
            ),
            columns=[
                "Low_prob_detections_0.1",
                "Low_prob_detections_0.5",
                "Low_prob_detections_0.75",
                "Low_prob_detections_0.95",
                "Low_prob_detections_0.99",
            ],
        )
        self.csv_df_combined = (
            pd.concat([self.csv_df_combined, p_brackets_results], axis=1)
            .reset_index(drop=True)
            .fillna(0)
        )

    def save_file(self):
        self.csv_df_combined = self.csv_df_combined.drop(
            self.col_headers_shifted, axis=1
        )
        self.csv_df_combined = self.csv_df_combined.drop(
            [
                "Compass_digit_shifted",
                "Direction_switch",
                "Switch_direction_value",
                "Compass_digit",
                "Compass_direction",
                "Angle_sin_cumsum",
                "Angle_cos_cumsum",
            ],
            axis=1,
        ).fillna(0)
        write_df(
            self.csv_df_combined.astype(np.float32), self.file_type, self.save_path
        )


# test = FishFeatureExtractor(config_path='/Users/simon/Desktop/envs/troubleshooting/naresh/project_folder/project_config.ini')
```
